# Development of Germline Progenitors in Larval Queen Honeybee ovaries

**DOI:** 10.1101/2024.04.22.590642

**Authors:** Georgia Cullen, Erin Delargy, Peter K. Dearden

## Abstract

Honeybees (*Apis mellifera*) are a keystone species for managed pollination and the production of hive products. Eusociality in honeybees leads to much of the reproduction in a hive driven by the queen. Queen bees have two large active ovaries that can produce large numbers of eggs if conditions are appropriate. These ovaries are also active throughout the long lives of these insects, up to 5 years in some cases.

Recent studies have indicated that the germline precursors of the adult honeybee queen ovary are organized into 8 cell clusters, joined together by a polyfusome; a cytoplasmic bridge. To understand the origin of these clusters, and trace the development of the honeybee queen ovary, we examined the cell types and regionalization of the developing larval and pupal queen ovaries.

We used established (*nanos* and *castor*), and novel (*odd skipped*) gene expression markers to determine regions of the developing ovary. Primordial germline cells develop in the honeybee embryo and are organized into ovary structures before the embryo hatches. The ovary is regionalized by Larval Stage 3 into terminal filaments and germaria. At this stage clusters of germline cells in the germaria are joined by fusomes and are dividing synchronously. The origin of the 8-cell clusters in the adult germarium is therefore during larval stages.

On emergence, the queen ovary has terminal filaments and germaria but has not yet developed any vitellaria, which are produced after the queen embarks on a nuptial flight. The lack of germaria, and the storing of germline progenitors as clusters, may be adaptions for queen bees to endure the metabolic demands of a nuptial flight, as well as rapidly lay large numbers of eggs to establish a hive.

## Introduction

*Apis mellifera* (Honeybees) are eusocial insects, characterised by cooperative brood care, overlapping generations and a division of reproductive labour (Wilson and Hölldobler, 2005). Individuals undergo enforced altruism (Ratnieks and Halenterä, 2009), becoming aggressive to workers with more developed ovaries (Visscher and Dukas, 1995) and killing pathogen-infected (Starks et al., 2000). The two female castes of honeybees, workers and queens, are exemplars of the polyphenism underlying the division of reproductive labour. Queen bee anatomy differs from workers, especially in the case of the abdomen, which is larger in queens, and contains two large active ovaries (Snodgrass, 1956). The queen’s ovaries are divided into up to two hundred ovarioles in each (Cridge et al., 2017), while workers have less than ten per ovary (Snodgrass, 1956). This polyphenism in ovary size is a result of differential feeding during larval developmental stages (Rachinsky et al., 1990). After eclosion, honeybee queens remain in the hive for a few days (Plate et al., 2019) to a week (Jackson and Robinson, 2018), before undertaking nuptial flights (Woyke, 1964). After mating, queens usually take a few days to begin laying (Kocher et al., 2008). The queen produces thousands of embryos that become either workers or drones depending on fertilization (Woyke, 1963). She is the only individual in a colony that can produce fertilized eggs, and thus female workers. Therefore, the queen’s ability to lay is crucial for long-term hive success (Gill and Hammond, 2011).

Honeybees have polytrophic meroistic ovaries (Gutzeit et al., 1993; Tanaka and Hartfelder, 2004). This ovary structure is defined by the presence of nurse cells, which provide nutrients and materials to the developing oocyte, and accompany it along the ovariole (Büning, 1993). In honeybees, the ovariole is divided into three major sections: the terminal filament, germarium and vitellarium (Cullen et al., 2023; Dearden, 2006). The terminal filament consists of flattened somatic cells, that appear to provide the progenitors of the somatic follicle cells of the ovary(Hartfelter and Steinbrück, 1997; Martins et al., 2011; Tanaka and Hartfelder, 2004). The germarium contains the first germ-line progenitors, which are clusters of 8 germline cells joined by a polyfusome (Cullen et al., 2023). These 8-cell clusters can divide synchronously at a rate that maintains the numbers of these clusters between laying seasons (Cullen et al., 2023).

The progenitor 8-cell clusters in the germarium will divide multiple times(Cullen et al., 2023; St Johnston, 2008) maturation of the oocyte, takes place in the vitellarium (Chapman, 1998). The oocyte must undergo meiosis, and accompanying nurse cells will divide and endoreplicate, producing polyploid nurse cells linked to each oocyte via the ring canals. The oocytes will then grow, and become provisioned with yolk until they are ready to be laid (Aamidor et al., 2022). The 8-cell cluster is a key point in reproduction as these are the primary store of germline cells in the ovary and the source of oocytes.

Despite the importance of honeybees to the economy and the environment (Khalifa et al., 2021; Tanaka and Hartfelder, 2004), honeybee populations are declining (Brosi et al., 2017; Yang et al., 2023). They experience a range of threats, such as parasites (Warner et al.), viruses (Abbo et al., 2017; Kang et al., 2016), and insecticide exposure (Manzoor and Pervez, 2022; Stuligross and Williams, 2021). Due to these threats, and others, the costs of beekeeping are increasing. A better understanding of how queen honeybees consistently produce large numbers of offspring, and how to optimise conditions to support that reproduction, will help reduce the effects these threats have on the global populations.

To understand how to best support honeybee reproduction we need to understand the dynamics of germline and oocyte production. Previous studies have shown that germline cells are specified late in honeybee embryogenesis (Dearden, 2006) and that the bulk of ovary growth occurs during larval and pupal stages(Chapman, 1998; Hartfelter and Steinbrück, 1997).

Here we set out to trace ovary development in queen honeybees. Understanding when and where ovary structure appears, how the ovarioles form, and, perhaps most importantly, where/when the 8-cell clusters form, is crucial information if we are to better support reproduction in queen bees and understand the evolution of the unusual structures of the honeybee ovary.

## Materials and methods

### Honeybees and staging

Honeybees were sourced from Otto’s Bees, a bee-keeping supplier based in Dunedin, Otago, New Zealand. Otto’s Bees keeps two stocks of bees, an ‘Italian’ strain, derived from a closed breeding programme maintained by Betta Bees Research Ltd, and a ‘Carniolan’ strain. Replicates of experiments were performed using unrelated hives.

Grafting of embryos into queen cells and queen cell raising was carried out as per)Cameron et al., (2013).

Embryos and larvae were staged according to Cridge et al., (2017). We investigated larval stages 2, 3 and 4. The larvae are in stage 5 for four days (Cridge et al., 2017) so we investigated ‘early’ and ‘late’ periods. We also split the pupal stage, also 5 days, into early and late stages.

The staging was performed by timing individuals from days post-laying, and based on morphology, as described below.

**Table 1:** Staging of individual honeybees used in these experiments based on Cridge et al., (2017).

| Stage | Days Post-Laying | Numbers of ovaries assayed across replicates |
| --- | --- | --- |
| Larval Stage 2 | 5 | 17 |
| Larval Stage 3 | 6 | 40 |
| Larval Stage 4 | 7 | 30 |
| Larval Stage 5 (Early) | 8-9 | 27 |
| Larval Stage 5 (Late) | 10-11 | 31 |
| Early Pupa | 12-13 | 39 |
| Late Pupa | 15-16 | 28 |
| Newly Emerged Queens | 17+ | 115 |

### Dissection and fixation

#### Embryos

Embryos were collected, fixed and dissected using a protocol modified from Osborne and Dearden, (2005), as described below. Embryos were collected from worker cells with a paintbrush into 1:1 heptane:4% formaldehyde in 1x PBS. Embryos were shaken overnight at moderate speed on a nutating mixer. After shaking, the lower phase was removed and replaced with 100% methanol at -20°C, this was then shaken vigorously for 3 minutes by hand. The solution was then removed, and embryos were washed 3 times in fresh -20°C 100% methanol. Embryos were then stored until dissections in 100% methanol at -20 °C.

Embryos and early larva were transferred to a thin-walled glass test tube; 100% methanol was replaced with 2 mL PTw (1x PBS + 0.1% Tween 20). Embryos were allowed to settle and then sonicated in a sonic cleaning bath for 5-10 seconds. Embryos were transferred to a microcentrifuge tube with 200 μg/mL of Pronase (Sigma) in PTw. After 5 minutes the embryos and larva were washed twice with 500μL of PTw. The embryos were transferred to a plastic petri dish with PTw and the outer membranes dissected away using sharp forceps. Dissected embryos were collected into a microcentrifuge tube with PTw. Embryos were then fixed again in 10% formaldehyde in PBS for 35 minutes. Embryos were then rinsed 6 times in 500μL of PTw.

### Collection of Larval and Pupal Stages

Larva and pupa were collected from queen cells. Larval samples were kept in the hive until necessary to continue brood care. Pupae were removed from the hive and kept in an incubator with at least 50% ROH at 34°C. Upon removal from queen cells, larvae and pupae were immediately dissected. Larval stages 2 and 3 were dissected by cutting a slit along the centre of the dorsal midline and loosening the internal organs with tweezers. The entire larva was then stained as below. For Larval stages 4 and 5, ovaries were dissected by cutting a slit along the centre of the dorsal midline and dissecting them from the larval body. For pupal stages, the head and thorax was removed from the abdomen. Then segments three and four were pulled apart and ovaries removed from the abdomen.

Newly emerged queens were always dissected at less than 12 hours old. Newly emerged queens were placed at 4°C until movement ceased, and then the heads were removed from the thorax before immediate dissection. Ovaries were dissected into 1x PBS under a dissection microscope and kept in 1x PBS on ice until fixation (less than 2 hours). Once ready for fixation, ovaries were fixed in 1:1 heptane:4% formaldehyde in 1x PBS for 12 (larval stages)-15 (pupal stages and newly emerged queens) minutes, then rinsed in 1x PTx (PBS + 0.1% Triton X-100) 6 times. All ovaries were used immediately in experiments.

### *In situ* Hybridization

#### Embryos

Honeybee embryos were stained via a modification of that used for *Nasonia* embryos in Taylor and Dearden, (2022). Embryos were pre-hybridised in 200μL of probe hybridisation buffer (2.4 M Urea, 5 × sodium chloride sodium citrate (SSC), 9 mM citric acid (pH 6.0), 0.1% Tween 20, 50 μg/mL heparin, 1 × Denhardt’s solution, 10% dextran sulfate) for at least 30minutes at 37°C and probe solution was prepared by adding 6μL of each 1 μM probe (Molecular Instruments) to 200 μL probe hybridisation buffer at 37°C. The pre-hybridisation solution was removed, and the probe solution was added. These samples were then incubated for 2 days at 37°C. Unbound probes were removed by 4 x 15-minute wash steps with 200μL of probe wash buffer (2.4 M Urea, 5 × SSC, 9 mM citric acid (pH 6.0), 0.1% Tween, 50 μg/mL heparin) at 37°C and 3 x 5-minute washes with 200μL 5X SSCT (5X SSC, 0.1% Tween 20) at room temperature. Embryos were pre-amplified in 200μL of amplification buffer (5X SSC, 0.1% Tween 20, 10% dextran sulfate) for at least 30 minutes at room temperature. Six μL of h1 and h2 hairpins (Molecular Instruments) were prepared separately, snap-cooled by heating to 95°C for 90 seconds then left to cool in the dark for 30 minutes. The hairpin solution was then prepared by adding all the hairpins to 200μL of amplification buffer at room temperature. The pre-amplification solution was then removed and replaced with the hairpin solution. Embryos were incubated for 2 days in the dark at room temperature. Unbound hairpins were then removed by 2 x 5-minute washes with 200μL of SSCT, 2 x 30-minute washes with 200μL of SSCT and 1 x 5-minute wash with 200μL of SSCT. SSCT was then removed and replaced with 500μL 70% glycerol. Embryos were then bridge-mounted for microscopy.

#### 2-5 Stage Larva/Pupal Stages

Hybridization chain reactions of larva and pupal ovaries were performed as described in Cullen et al., (2023), with probe concentrations of: 2μL of 1 μM probes (*nanos, castor, odd skipped*). Hairpins were changed between replicates (-488,-546,-647), to ensure no signal bias in specific wavelengths of light.

### Immunohistochemistry

Immunohistochemistry, phalloidin and DAPI staining was performed as described in Cullen et al., (2023), with primary antibody for dividing cells ((1:200 α-mouse-ph3 (Anti-Histone H3 (phospho S10) antibody [abcam 14955]).

### Microscopy

Confocal microscopy was performed on an Olympus FV3000 confocal microscope. Ovaries were serially optically sectioned with the number of slices/depth optimized using FV3000 software. Images were produced using Icy(de Chaumont et al., 2012) or FIJI (Schindelin et al., 2012) and images trimmed for ease of visualisation. All raw data images and media are available at Zenodo (DOI 10.5281/zenodo.10972330).

### Phylogenetics

Proteins with similarities to odd skipped-like proteins were identified using blastp (Altschul et al., 1990), and aligned using Clustal (Thompson et al., 1994). Maximum likelihood phylogenetic analysis was carried out using RAxML (Stamatakis, 2014).

## Results

### Timing of Development of Honeybee Queen Ovary Morphology

The precursors to honeybee ovaries form in the embryo as two lines of cells in the most dorsal regions of stage 9 embryos, between abdominal segments 3 and 6 (Dearden, 2006). These consolidate into two arches of cells underlying the dorsal surface of just-hatched (L1) larval honeybees (Dearden, 2006). Considerable growth and differentiation occurs during larval and adult stages.

Stage 1 and 2 ovaries are difficult to visualize and dissect out of the larval body, and do not stain well if kept as whole mounts. L2 larval ovaries lie on either side of the midline between abdominal segments 3 and 6. The ovaries are small and transparent. The cells of the ovary are linked with trachea and buried in the fat layer of the L2 larva (Figure 1(A)).

**Figure 1:**
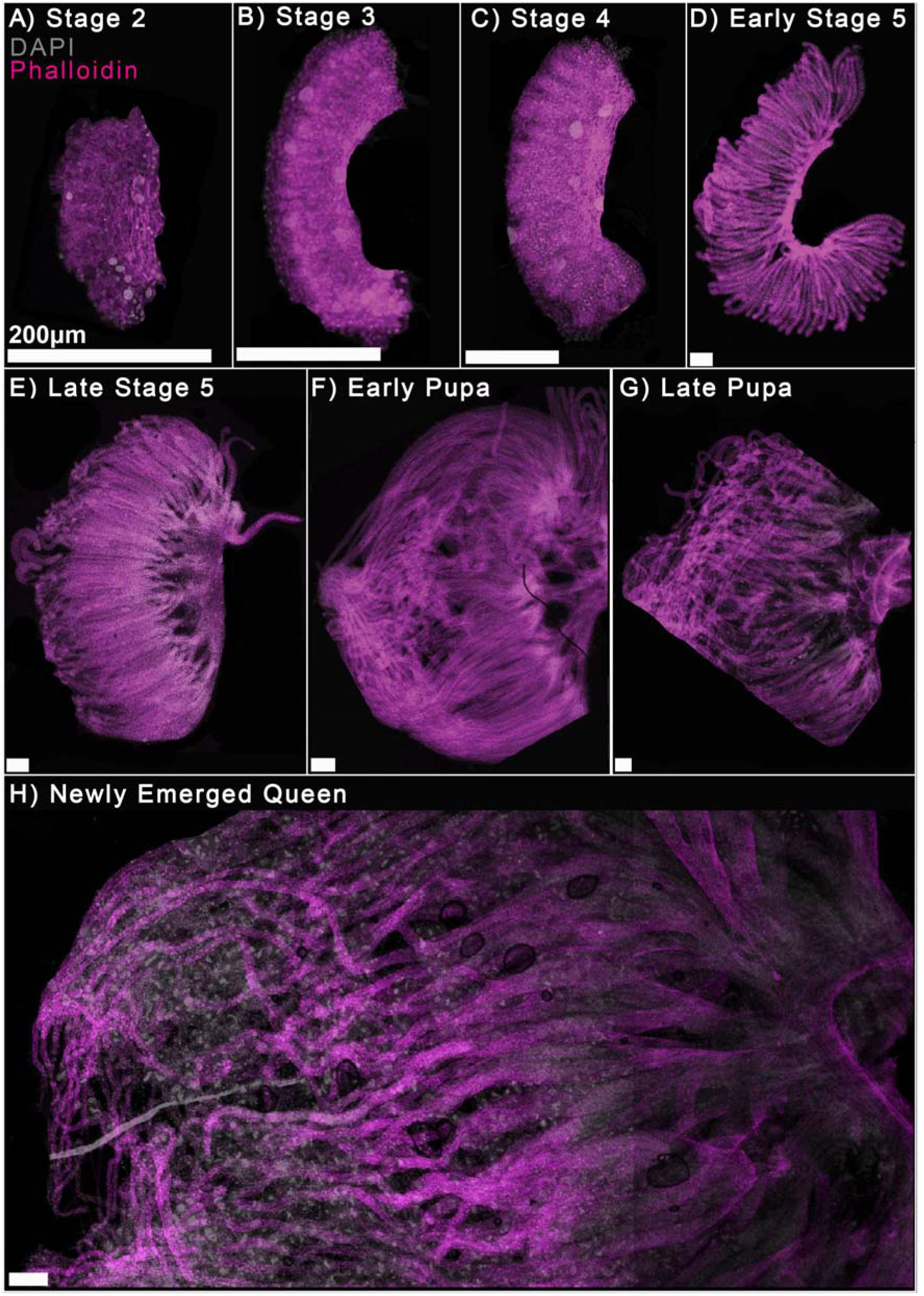
A developmental series Honeybee (*Apis mellifera lingustica*) queen ovaries. The ovaries are oriented with the developing terminal filaments to the left, forming oviduct to the right. Ovaries stained for DNA with DAPI (grey) and cortical actin with phalloidin (magenta). Images are organized by stage: A) larval stage 3, B) larval stage 4, C) early larval stage 5, D) late larval stage 5, E) early pupa, F) late pupa and G) newly emerged queen. Scale bars are 200 µm.

L3 queen larval ovaries have contracted somewhat and are now found between abdominal segments 4 and 5 on either side of the dorsal line (data not shown). When stained with DAPI and phalloidin, the cells are organized into ‘fingers’ of tissue that resemble small future ovarioles, although they are not separated from each other (Figure 1(B)).

From this stage onward, the ‘fingers’ of tissue elongate, and, by early L5, separate into individual ovarioles. The ovarioles lengthen through late larval stage 5, with connective tissue forming to connect the posterior of all the ovarioles (Figure 1(E-G)). This is likely tissue that will go on to be, or connect to, calices (Kozii et al., 2022). Calices provide the transfer of eggs between the ovaries and the lateral oviduct (Kozii et al., 2022). This region connects both lateral oviducts from either ovariole with the spermatheca (median oviduct)(Kozii et al., 2022).

Growth continues in the pupal stages, with terminal filaments and germaria present. On hatching, the ovary does not appear to contain nurse cells or differentiated oocytes.

### Regionalisation of the ovary during development

In order to understand the regions of the developing honeybee ovaries, we used markers of specific cell types to identify each region as it develops. In adults, ovary cells that express *castor* (*cas*) are somatic (Cullen et al., 2023) and so we used this gene as a potential marker of somatic cells in the ovary. Germline cells express *nanos* (*nos*) and *vasa* (*vas*)(Dearden, 2006) from late embryonic stages through to adults (Cullen et al., 2023), allowing us to use these as markers of germline fate. RNA expression from the *nos* gene was used in these experiments to mark germline cells

The *odd skipped (odd)* gene encodes a zinc finger transcription factor that in *Drosophila melanogaster* acts as a pair-rule gene (Coulter et al., 1990). *Odd* is a member of a family of similar transcription factors encoded by the *brother of odd with entrails limited* (*bowl*) and *sister of odd* (*sob*) genes (Hart et al., 1996). To determine the most likely orthologue of *odd* in honeybees, we identified homologues of all these genes using blastp (Altschul et al., 1990), and then carried out maximum likelihood phylogenetics. Honeybee XP_001120949 clusters with odd proteins from other holometabolous insects against sob and bowl proteins (Supplementary Figure 1). We thus designate the gene encoding this protein, honeybee *odd skipped* (*odd*).

In *Drosophila*, RNA from *odd* is expressed in a variety of cell types, but not germline or ovary cells (Coulter et al., 1990; Ward and Coulter, 2000). In experiments to examine the expression of genes involved in segmentation in honeybees (data not shown), we identified that honeybee *odd skipped* is expressed during segmentation, but is also expressed in the same presumptive ovary region as *nos* and *vas* RNA in late embryos (Figure 2). This region, located in the most dorsal regions of the germband, between segments 3 and 6, is where primordial germline cells can first be detected via expression of *nos* and *vas* RNA(Dearden, 2006).

**Figure 2:**
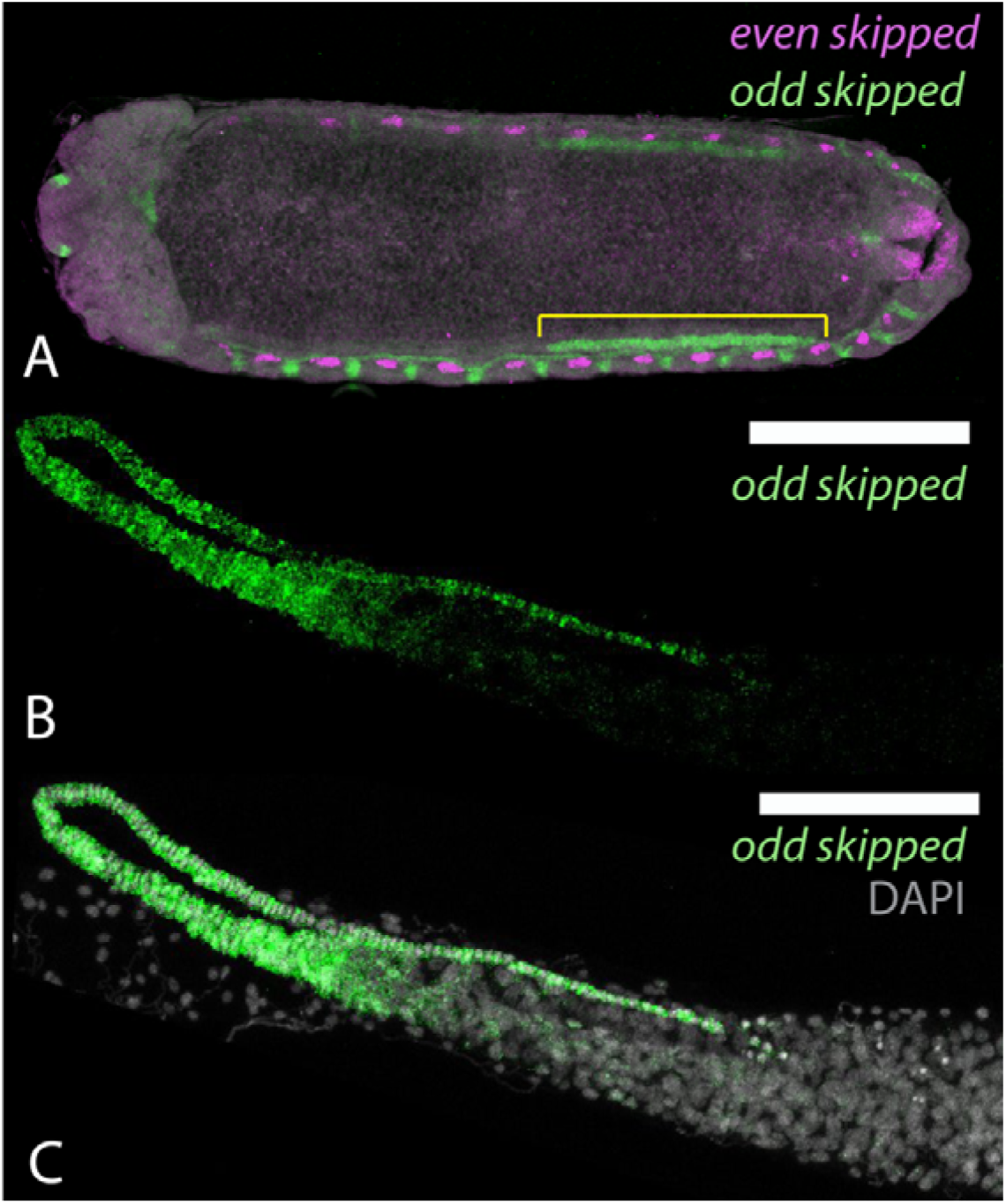
Expression of honeybee *odd skipped* in embryos and adult ovaries. A) Stage 8 honeybee embryo (Cridge et al., 2017) stained for *even skipped* (magenta) for reference (Wilson and Dearden, 2012) and *odd skipped* (green)(Supplemental Figure 1). *Odd skipped* RNA is expressed in clusters of cells interspersed with even skipped expression along the body wall of the embryos, and in a line of cells stretching from abdominal segment 3 to abdominal segment 6 in the most dorsal region of the germ band (marked by yellow line). B and C) Expression of *odd* (green) in terminal filament regions of an adult queen ovary. RNA expression appears in all the cells of the terminal filament (in this specimen folded over the germarium of the ovary), with expression reducing at the boundary of the germarium. DNA stained with DAPI, grey in C.

To attempt to determine what cells *odd* might be marking in the honeybee embryonic ovary, we investigated where *odd* is expressed in the adult queen ovary. RNA from *odd* is expressed in the cells of the terminal filament (Figure 2). It is detected in the cells around the boundary of the terminal filament and the germarium, but absent in the follicle cells surrounding the germarium (Supplementary Figure 2). These cells are thought to be the descendants of terminal filament cells (Dobens and Raftery, 2000). This encouraged us to use *odd* as a marker for presumptive terminal filament cells during ovary development.

At Larval stage 3, *odd, cas* and *nos* RNA expression can be used to identify the cell-type structure of the developing ovary (Figure 3). The ovary is a curved structure with the outside of the curve oriented to the midline of the larva. Many of the cells of the ovary appear to express *castor* RNA, especially the trachea that infiltrate the tissue. Expression of *odd* RNA appears in clusters of cells on the outside of the curved ovary. RNA from *nos* appears in groups of cells juxtaposed to *odd* expressing cells. Close analysis of these expression patterns (Figure 3F, G and H) show *odd* RNA expression at the end of the small fingers of tissue in the ovary at this stage. RNA from *nos* is expressed in a band of cells abutting the *odd* domain further down the structure. Cells do not co-express *odd* and *nos* RNA. This relationship is very similar to the final pattern of gene expression of *odd* and *nos*, with *odd* RNA being expressed in the terminal filament, and its expression ceasing at the boundary of the germarium, where groups of *nos* RNA-expressing cells first appear in the ovary. These expression patterns imply that the finger structures are presumptive ovarioles, and these are initially separated into terminal filament and germline cells.

**Figure 3:**
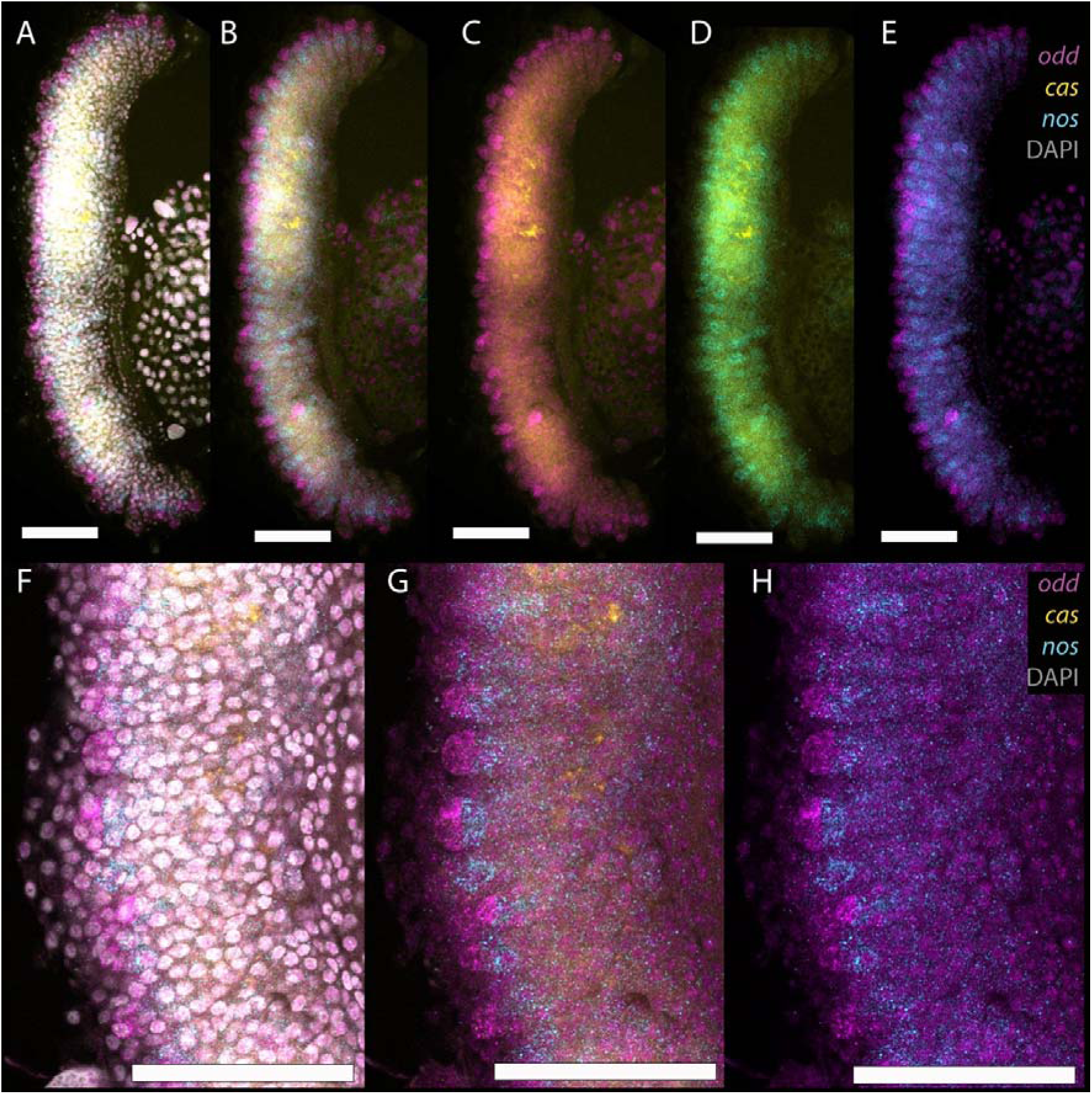
Expression of RNA from *cas* (yellow), *nos* (cyan) and *odd* (magenta) in L3 larval ovaries. A) Ovary stained with DAPI (grey), *cas*, *nos* and *odd*. B) *cas*, *nos* and *odd*. C) *cas* and *odd*. D) *cas* and *nos*. E) *nos* and *odd*. F, G and H) close up of expression of F) DAPI, *cas*, *nos* and *odd*. G) *cas*, *nos* and *odd* and H) *nos* and *odd*. Expression of *odd* RNA (magenta) can be seen at the end of the small ‘fingers’, with *nos* RNA expression in a cluster of cells abutting the *odd* domain. Scale bars indicate 100 μm.

Larval stage 4 ovaries show a very similar arrangement as those at L3. Though the ‘fingers’ are slightly longer. By early larval stage 5, however, the ‘fingers’ have separated into ovarioles, with the terminal tips of the ovarioles expressing *odd* and *cas* RNA, *nos* RNA being expressed by clusters of cells further down each ovariole, and *cas* RNA expressing cells surrounding these. No cells co-express *cas* and *nos* RNA.

As the larval ovary extends during L5, the arrangement of cells seen in early L5 is maintained but extended. *Odd* and *cas* RNA-expressing cells make up the developing terminal filament. Moving along each ovariole, in the region in which clusters of *nos* RNA expressing cells appear, *odd* RNA-expressing cells disappear, with cells surrounding the *nos* RNA expressing cells expressing only *cas* RNA. These structures and gene expression patterns remain throughout larval and pupal development as the ovary grows. By analogy to the adult form, the cells of the terminal filament and the germline are defined in the embryo, they become organized by L3, and then divide and grow to produce the structures of the terminal filament and germarium in the adult ovary. There is no evidence for a vitellarium in larval or pupal ovaries.

These *in situ* hybridization data indicate that there are germline cells in groups at L3. However, it does not tell us whether these germline cells are linked via cytoplasmic bridges, and divide synchronously, as in adult germline cell clusters.

### Formation of intercellular bridges

Having described the regionalisation of the ovary into terminal filaments and germaria, we next investigated when and where the germline cells form into the 8-cell clusters that comprise the germline in adult queens (Cullen et al., 2023). These clusters have two key characteristics. The first is that they are joined by a polyfusome, a cytoplasmic bridge that links the 8 cells together (Cullen et al., 2023). The second characteristic is that they divide synchronously (Cullen et al., 2023). To determine when 8 cell clusters begin to form in the ovary, we stained for polyfusomes with phalloidin (Cullen et al., 2023), and for dividing cells with antibodies directed to phosphorylated histone H3 (Hans and Dimitrov, 2001).

Fusomes and polyfusomes are cytoplasmic intercellular bridges between cells forming cystocytes and germline cell clusters (Ventelä, 2006). In *Drosophila melanogaster,* fusome precursors are called spectrosomes (Deng and Lin, 2001). The cystoblast spectrosome grows into a fusome, a distinct region of the cytoplasm, which bridges two germline cells(King, 1979), as germline cysts form (de Cuevas and Spradling, 1998). Polyfusomes, in *Drosophila*, are assembled from further joining of fusomes creating branched structures that can link together the cytoplasm of several cells (King, 1979; Pritsch and Büning, 1989; Trauner and Büning, 2007). The appearance of polyfusomes during development implies the formation of germline cysts (Cullen et al., 2023).

Phalloidin staining should mark spectrosomes, fusomes, polyfusomes and ring canals. In the earliest larval stage ovaries we can image with confidence, larval stage 2, we cannot find accumulations of phalloidin staining consistent with any of these structures. By larval stage 3, however, phalloidin staining can be seen staining concentrations of material that appear to link neighbouring cells (Figure 5B), reminiscent of *Drosophila* fusomes. These potential honeybee larval stage 3 fusomes are 5-7.6 μm wide comparable to *Drosophila* fusomes, (6-10 μm wide during cystoblast to 2-cell cyst stages)(de Cuevas and Spradling, 1998). The similarity in size, staining and shape implies the presence of fusomes at larval stage 3, and that some of the germline cells have formed 2 or 4-cell cysts.

**Figure 4.**
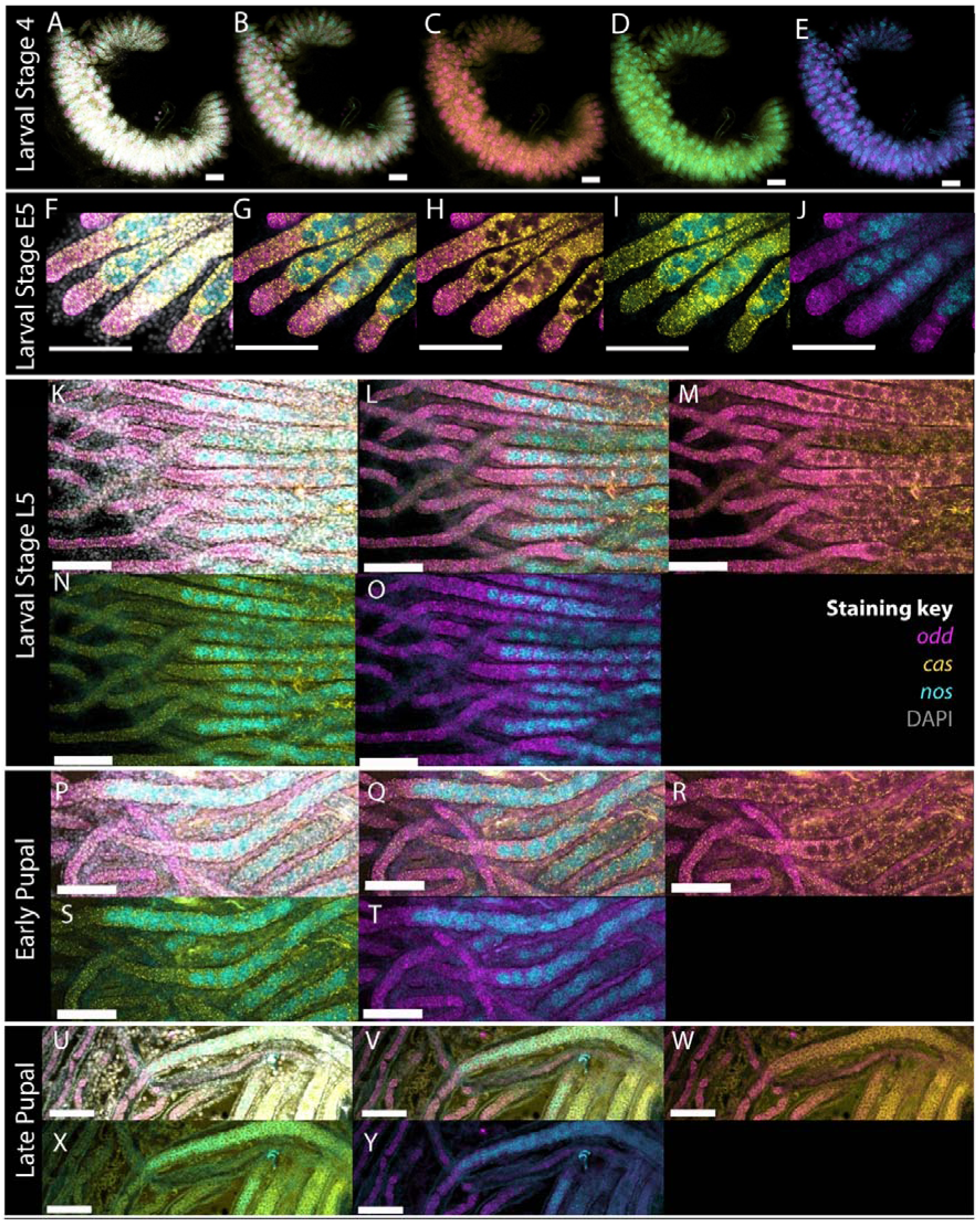
Expression of *cas* (yellow), *odd* (magenta) and *nos* (cyan) throughout larval and pupal development of queen honeybees. DNA stained with DAPI is shown in grey. A-E) L4 ovaries, with staining for DAPI, *cas*, *odd* and *nos* (A), *cas*, *odd* and *nos* (B), *cas* and *odd* (C), *cas* and *nos* (D) and *odd* and *nos* (E). Other stage ovaries are stained with the same genes and displayed in the same colours. Scale bars are 100 µm.

**Figure 5.**
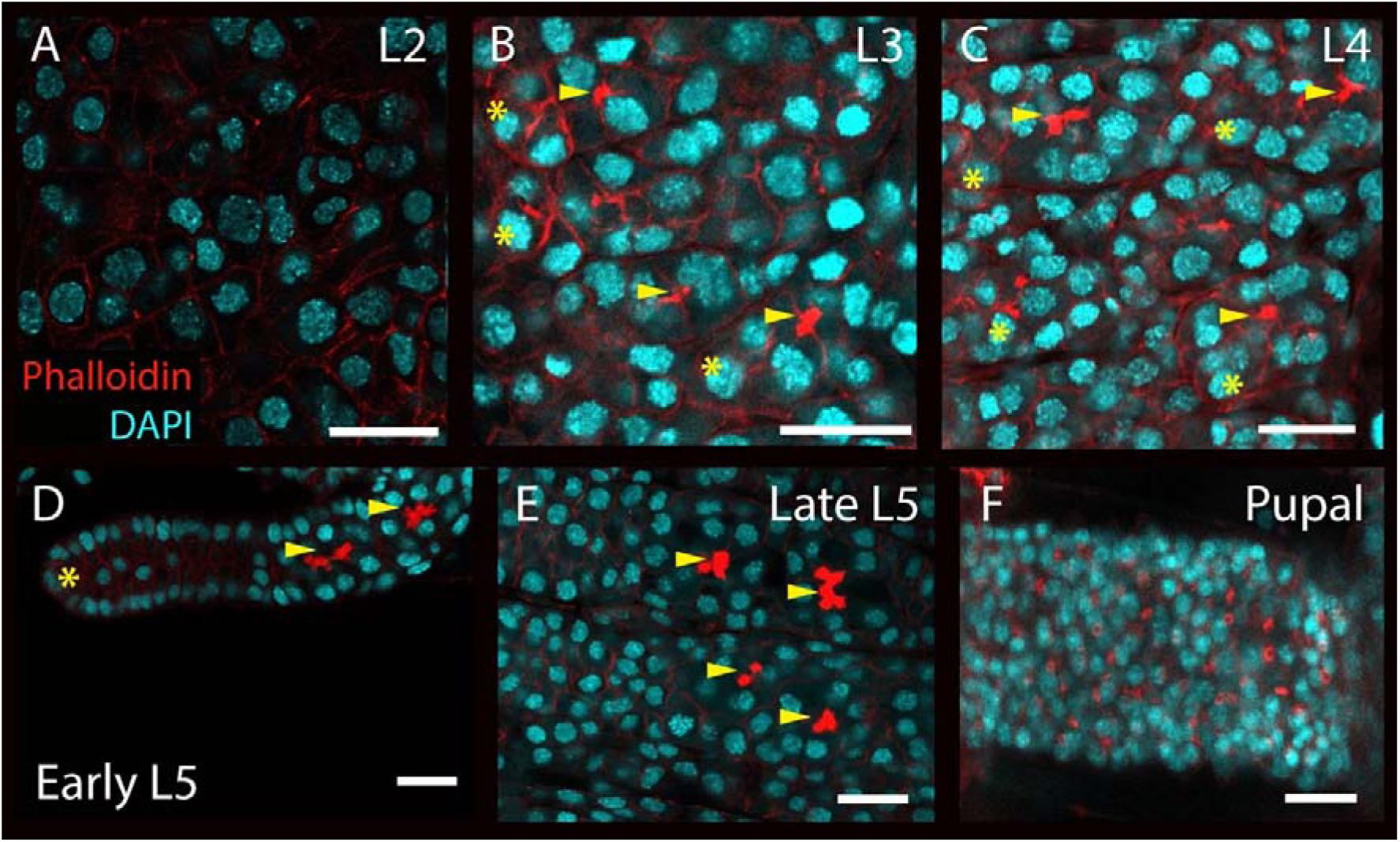
Phalloidin and DAPI stained larval and pupal ovaries focusing on the appearance of cytoplasmic bridges. A) Stage L2 ovary. Phalloidin stains cortical actin around cells, but there is no sign of conglomerations of staining implying the formation of fusomes. B) Stage L3, structures staining strongly with phalloidin can be seen bridging some neighbouring cells (marked with yellow arrowheads). These structures are generally to be found in cells adjacent to cells at the tips of the ‘finger’ structures in the ovary (marked where visible with yellow asterisks). C) Early stage L4 ovariole, strongly staining polyfusome-like structures (marked with yellow arrowheads) can be seen several 10s of micrometres away from the terminal structures of the ovariole. D) Late stage L4 ovaries; rows of phallodin-stained polyfusome-like structures can be seen in each ovariole. E) Pupal ovaries, polyfusomes structures can be seen as in late L4 ovaries (data not shown), and in more posterior regions, polyfusomes have disappeared and ring canals are now visible (phalloidin staining rings in E). Scale bars are 20 µm.

Polyfusomes, similar in size and shape to those in adult ovaries, are present from larval stage 4 (Figure 5C, D and E). Ring canals, which are present in the posterior germarium of adult queen honeybee ovaries, are not present in the ovary until late pupal stages (Figure 5F).

Phalloidin staining implies that fusomes, and thus the development of cysts, appear from larval stage 3. If these are the developing germline cell clusters, we should be able to detect synchronously dividing cells at the same stages.

### Synchronously dividing cells

Germline cell clusters in adult honeybee ovaries are joined by polyfusomes and divide synchronously (Cullen et al., 2023). Synchronously dividing cells, especially if associated with polyfusomes, in the developing ovary may be signs of clusters forming. As noted (Figure 1), the ovaries grow significantly during larval and pupal stages. This growth is associated with large amounts of cell divisions in all tissues of the developing ovary (Figure 6) making identifying synchronously dividing cells challenging.

**Figure 6.**
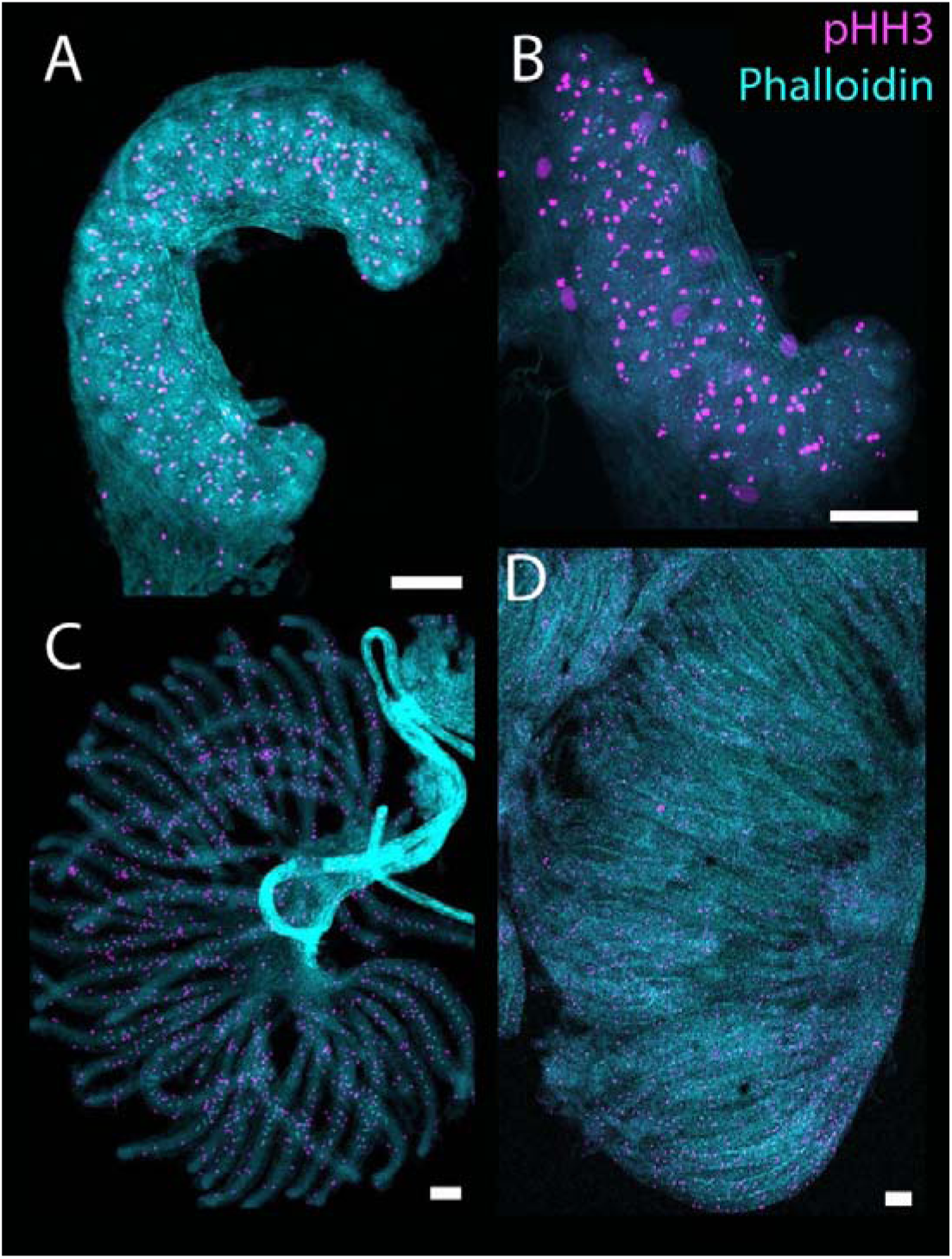
Immunohistochemistry for PHH3 in larval and pupal ovaries to detect dividing cells. Ovaries are stained with PHH3 (magenta) and phalloidin (cyan) to show where dividing cells appear in the developing ovary. As the ovary grows, many cells in the structure divide. A) L3 ovary. B) L4 Ovary. C) Early L5 ovary, note the polyfusomes lined up in each ovariole stained with phalloidin (cyan) D) Late L5 ovary. Ovaries are orientated with the terminal filament (anterior) to the left. Scale bars are 100 µm.

Synchronously dividing cells can be detected during larval ovary development. At larval stages 3 (Figure 7A and Supplemental Video 1) and 4 (Figure 7B and Supplemental Video 2), there are clusters of simultaneously dividing cells in the region of *nos* positive cells in the ovary. At larval stages 3 and 4, these are clusters of 4 cells, while at larval stage 5 (Figure 7C and Supplemental Video 3) these are clusters of 8 dividing cells. These dividing cells, while in the same regions of polyfusomes, have very faint, or difficult to see polyfusomes in the same region. When the germline cell clusters in adult ovaries divide, the phalloidin staining of the polyfusomes joining them is reduced, making them hard to discern (Cullen et al., 2023).

**Figure 7.**
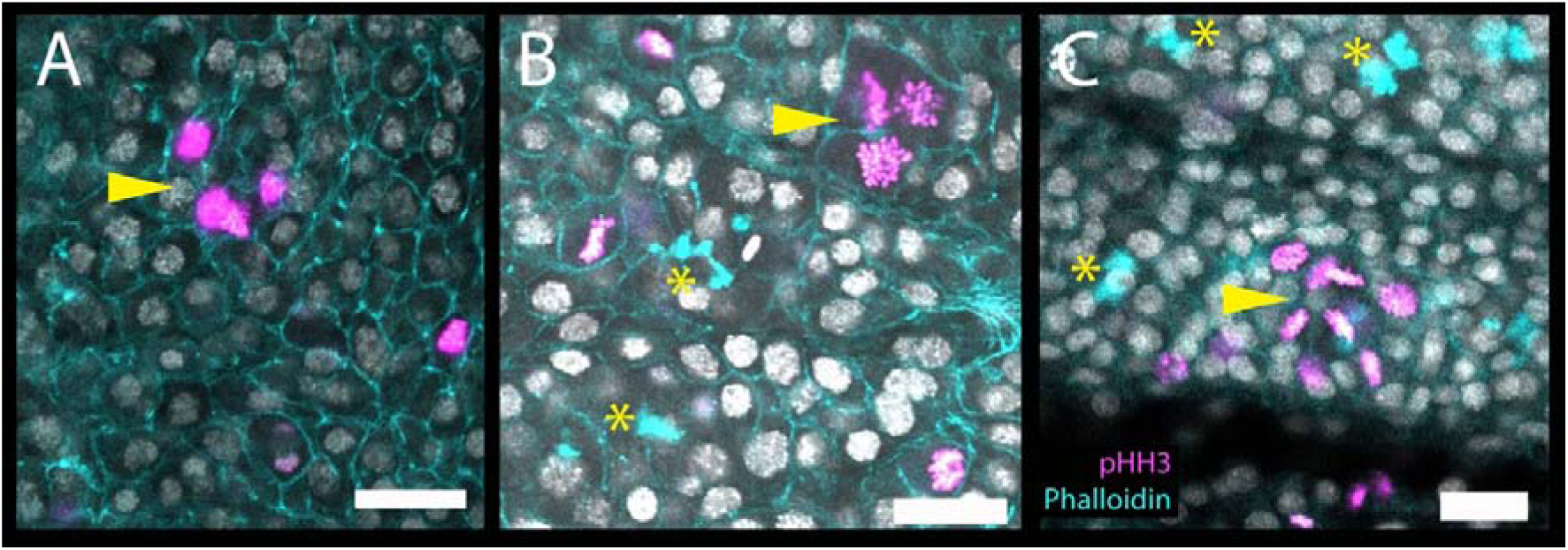
Clusters of synchronously dividing cells during larval ovary development. Ovaries are stained for PHH3 (magenta) and cortical actin using phalloidin (cyan). See also supplemental videos 1,2 and 3. A) Larval stage 3 ovary showing a cluster of 4 synchronously dividing cells (arrowhead) B) L4 stage ovary showing polyfusomes (asterisks) and a cluster of 4 dividing cells (arrowhead). C) Early L5 ovary showing two ovarioles, both with polyfusomes (asterisks) and a cluster of 8 synchronously dividing cells (arrowhead). Scale bars are 20 µm.

The appearance of fusomes, and then polyfusomes, as well as clusters of synchronously dividing cells, implies that 8-cell germline clusters in the adult ovary begin to form in larval stages 2 and 3, with the advent of cells joined by fusomes, which then, in larval stage 4, produce clusters of 8 cells joined together by polyfusomes. This clustered organisation of the germline in larval development remains the core structure of the germline in adult queen honeybees.

### Pupal growth of the ovary

As noted previously, the pupal ovary does not contain the vitellarium region of the ovary. In adults, the vitellarium contains meiosis and cysts of germ-line nurse cells and oocytes. Oocytes are provisioned and enlarge in the vitellarium, finally producing mature oocytes for laying. To determine when the vitellarium begins to form, we followed the expression of *cas*, *nos* and *odd* through late pupal development and into newly-emerged queens as a way to understand the further differentiation of the ovary (Figure 8).

**Figure 8.**
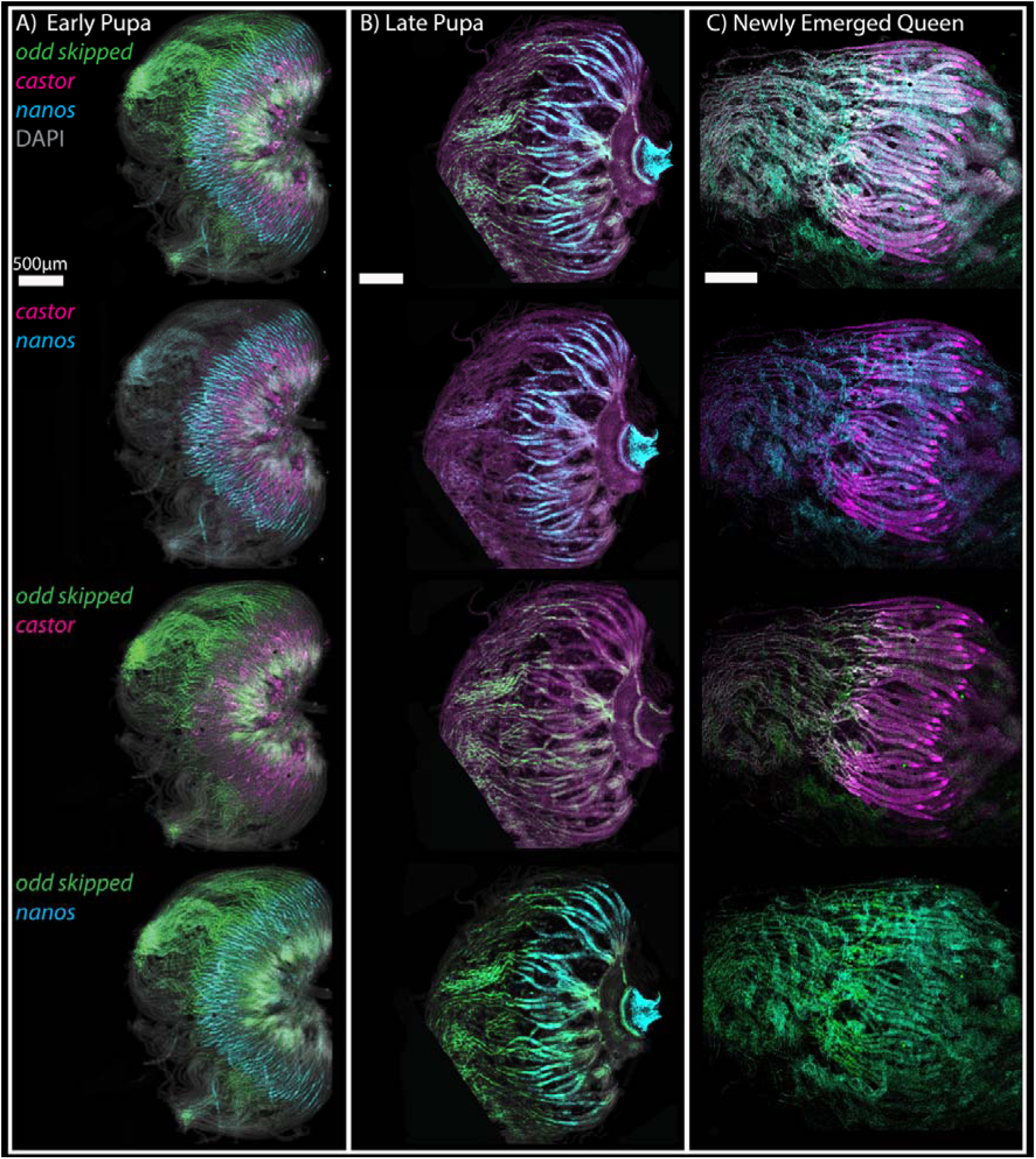
Later development of pupal and newly-emerged queen ovaries. Ovaries are stained for DAPI (grey), *odd* (green), *cas* (magenta) and *nos* (cyan). A) Early pupal ovaries. B) Late Pupal ovaries. C) Ovaries from a newly-emerged queen. Ovaries are orientated with the terminal filament (anterior) to the left. Scale bars are 500 µm.

The patterns of gene expression noted during larval development continue into late pupal and ovaries of newly-emerged queens. *Odd* expression marks the terminal filaments, with all terminal filament cells expressing *odd* RNA, until the anterior of the germarium, where clusters of *nos* expressing germline cell clusters reside. Expression of *cas* is restricted to somatic cells of both the terminal filament and the germarium. Expression of *nos* RNA is limited to the germarium.

Even in newly-emerged queen ovaries, there is little development of the vitellarium, supporting the idea that the ovary undergoes significant changes in structure and gene expression after mating(Kocher et al., 2008). Expression of *cas* RNA in newly-emerged queens marks a small triangular structure at the end of each germarium (Figure 8C), perhaps a sign of the beginnings of differentiation of the vitellarium.

While the vitellarium does not appear to form until after mating(Kocher et al., 2008), we wanted to know if the division of germline cell clusters is occurring in newly-emerged queens. By staining newly-emerged queen ovaries with an antibody against PHH3 we can see 8 cell clusters undergoing duplication in the germarium (Figure 9).

**Figure 9:**
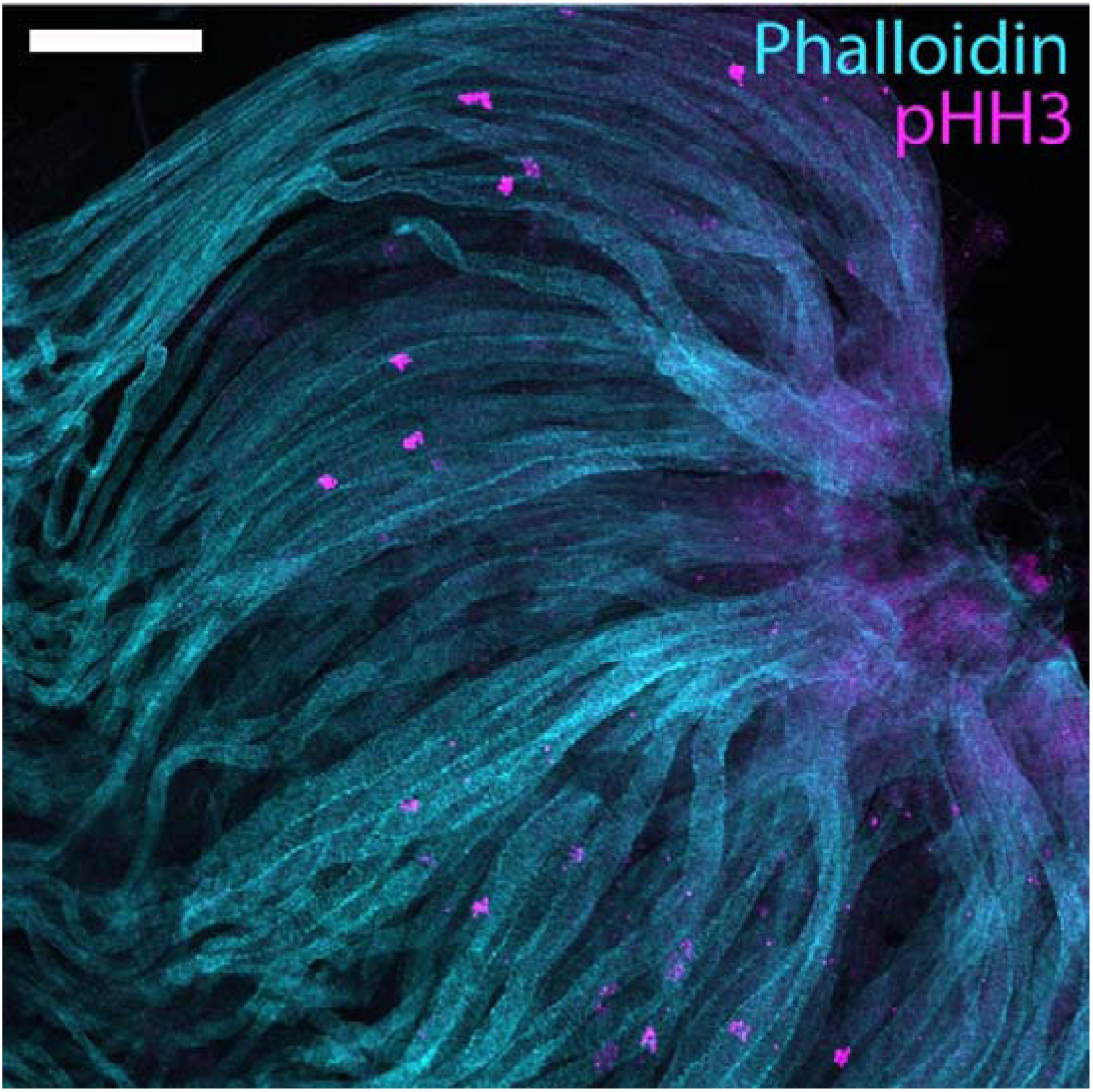
Ovary of a newly emerged queen honeybee stained for pHH3 (magenta) and cortical actin (cyan). Clusters of synchronously dividing cells can be seen in each ovariole.

## Discussion

Honeybee queens have been reported as laying ∼500-3000 (Allen, 1955; Avni et al., 2014) eggs per day. This egg-laying occurs in bursts, with permissive environmental conditions, and space in the hive required to allow these high rates of reproduction (Fine et al., 2018; Shehata et al., 1981). While the rate of laying is subject to environmental factors, e.g. nutrition and climate (Avni et al., 2014), structures that might affect egg production, such as ovariole number, can differ between queens (Jackson et al., 2011; Tarpy et al., 2000). It is also possible that the number, and quality, of germ-cell clusters can be manipulated during development, alongside ovariole number.

Understanding the development of the larval ovaries in bees required suitable markers. While *nos* and *cas* have been used previously, we serendipitouslydiscovered that *odd* is expressed from early embryonic stages, in the cells of the developing terminal filament. This expression does not occur in *Drosophila*, and we have, as yet no function for *odd* in the terminal filament. As a marker of terminal filament cells and development, *odd* expression demonstrates in bees, at least, that the terminal filament is an early developing component of the ovary; perhaps providing the somatic precursors to each ovariole. The initial ovary is thus a collection of terminal filament cells as somatic precursors to the ovary, and germline cells, all of which form in the embryo. Terminal filament cell number determines ovariole number in *Drosophila melanogaster* (Sarikaya et al., 2012), and co-varies with ovariole number in other *Drosophila* species (Green II and Extavour, 2012). Given the unusually large number of ovarioles in *Apis mellifera*, differing from most other Hymenoptera (Hartfelder et al., 2018), *odd* may provide a key marker in understanding the genetic mechanism underlying the evolutionary increase in ovariole number. Comparing the numbers of *odd* RNA-expressing cells in late embryogenesis between honeybees and Hymenoptera with smaller numbers of ovarioles and determining how they are specified may help us understand how, and why, queen honeybees produce so many ovarioles.

Tracing the regionalization of the honeybee ovary during larval development indicates that the ovariole structure is determined by larval stage 3, with terminal filament and germline cells residing in different domains by this early stage (Figure 10). Later larval and pupal development involves the expansion and elongation of these domains to produce an ovary in newly-emerged queens that is made up of an extensive terminal filament, and a germarium, but with no sign of the vitellarium. It is unclear if meiosis has begun in these ovaries as we have no markers to date that identify cells in meiosis in honeybees.

**Figure 10.**
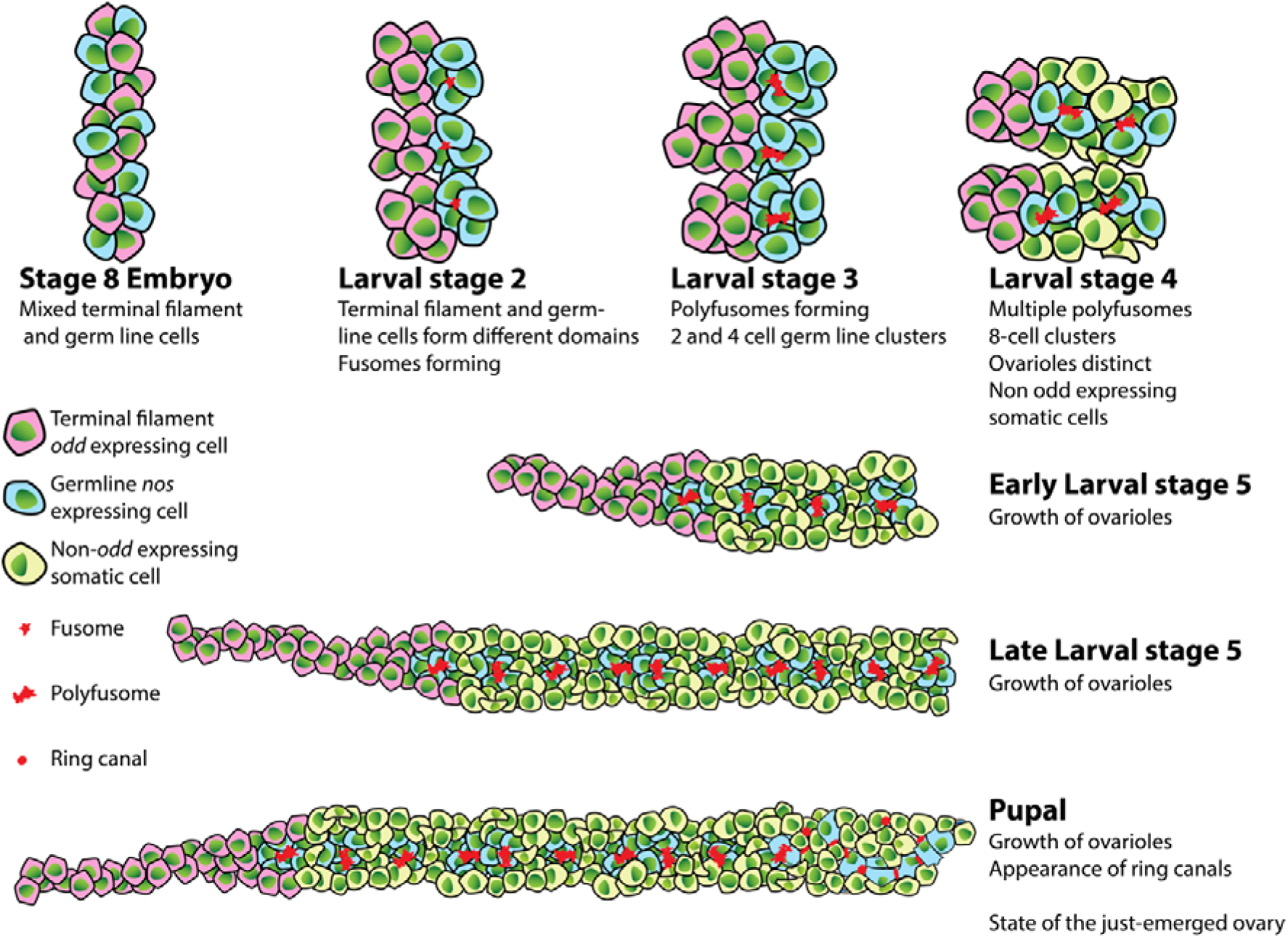
Cartoon of larval and pupal ovary development in honeybee queens indicating key events in the production of adult ovaries

The key feature of the honeybee queen germline is that germline precursors are organized into 8 cell clusters, joined by a polyfusome (Cullen et al., 2023). These structures reside in the germarium, where they divide at a rate that can replace these germline cell precursors over the years of a queen’s life (Cullen et al., 2023). We see evidence of the formation of these clusters beginning in Larval Stage 2, where we first see fusomes linking two cells in the germline regions of the ovarioles together, and stage 3 where we find synchronously dividing 4 cell clusters and polyfusomes. By early stage 5, there are numerous polyfusomes in the germarium, and clusters of 8 cells dividing synchronously (Figure 10). The germline cell clusters crucial to fertility in the adult queen honeybee thus form during early larval development and are in place in the germarium before the emergence of virgin queens.

The honeybee queen ovary is immature on emergence, as the vitellarium, where oocytes are specified and resourced, has not yet formed. Germline cell clusters are dividing at this time, but it is unclear if these clusters are being stored ready for reproduction to begin, or apoptosed. At this stage, after a few weeks in the hive (Koeniger, 1986), the virgin queens go on a mating flight, mating with multiple males on the wing, and then return to the hive. Egg laying begins 2-3 days (Koeniger, 1986) after mating. Changes in gene expression and morphology of the ovary occur after mating (Kocher et al., 2008), which produces an active vitellarium, and active egg laying.

Queen bees are constrained somewhat by their social biology. They need to be able to lay bursts of eggs when conditions are appropriate, reaching remarkable levels of egg laying (Harbo, 1986), and thus need large and active ovaries. Despite this, virgin queen bees also need to go on a metabolically challenging nuptial flight early in life. It seems likely that the immature ovaries in newly emerged queen bees reflect a trade-off in resource need for flight and reproduction. By suppressing, or delaying, the development and provisioning of eggs before mating, queens have the resources for an expensive nuptial flight. Trade-offs between reproduction and flight have been identified in many insect species (for examples see Guerra, (2011)). After repressing reproduction for a nuptial flight, queens need to rapidly activate their ovaries and begin laying large numbers of eggs to populate the hive with workers. The number of ovarioles in bees is unusual, especially for Hymenoptera (Church et al., 2021; Khila and Abouheif, 2010; Kugler et al., 1976; Martins and Serrão, 2004), a testament to the need to rapidly lay large numbers of eggs in ideal situations. Perhaps the large number of ovarioles in honeybee queens, and the 8-cell cluster form of the germline in the ovary (Cullen et al., 2023), are adaptations to allow the rapid activation and high rates of reproduction in young honeybee queens.

## Supplementary files

Supplementary file 1: Phylogenetics of odd-skipped proteins from holometabolous insects

Supplementary movie 1: Progressive focus-through of a larval stage 3 ovary stained for phalloidin (cyan) and pHH3 (magenta) to detect dividing cells. The focus passes through a 4-cell cluster of cells dividing synchronously.

Supplementary movie 2: Progressive focus-through of a larval stage 4 ovary stained for phalloidin (cyan) and pHH3 (magenta) to detect dividing cells. The focus passes through a 4-cell cluster of cells dividing synchronously.

Supplementary movie 3: Progressive focus-through of a larval stage 5 ovary stained for phalloidin (cyan) and pHH3 (magenta) to detect dividing cells. The focus passes through a 8-cell cluster of cells dividing synchronously.

## Data availability

The Authors affirm that raw data files for all figures are available on Zenodo ((DOI 10.5281/zenodo.10972330)

## Acknowledgements

The authors would like to that Dr Otto Hink for the provision of bees and grafting services and Amanda Austin for honeybee embryo staining. We thank Petra Dearden for critical review of the manuscript.

## Author contributions

G.C.: Experimental procedures, honeybee dissection, staining, imaging and image analysis, figure production, manuscript drafting. E.D.: Honeybee embryo collection and staining, imaging and image analysis. P.K.D.: Study conception, funding, image analysis, figure production, manuscript drafting, supervision.

---

**Supplemental Figure 1.**
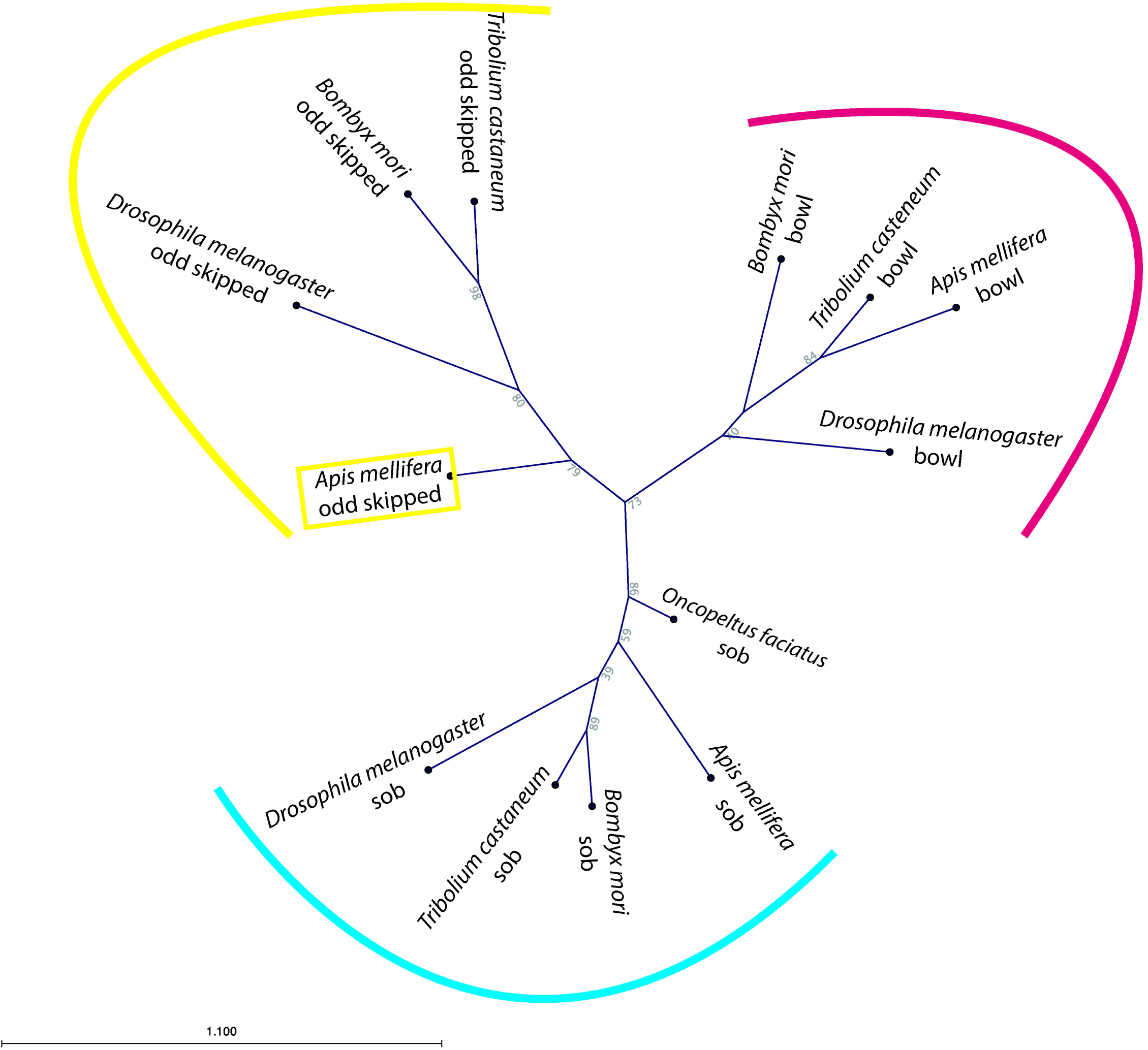
Maximum-likelihood phylogram of representative insect odd skipped like proteins. *Apis mellifera* odd skipped (yellow box) is in a clade with odd skipped from other insects to the exclusion of sob and bowl proteins. Sequence identifiers are; *Drosophila melanogaster* odd, FBpp0077246; *Drosophila melanogaster* bowl, FBpp0077180; *Drosophila melanogaster* sob, FBpp0077247; *Apis mellifera* odd, XP_001120949; *Apis mellifera* sob, XP_026300496; *Apis mellifera* bowl, XP_026300446.1; *Tribolium castaneum* odd, XP_008196754.1; *Tribolium castaneum* sob, XP_008196753.1; *Tribolium castaneum* bowl, XP_008196755.1; *Bombyxmori* odd, XP_037877356.1; *Bombyxmori* sob, XP_012550570.3; *Bombyxmori* bowl, XP_004927088.3; *Oncopeltus faciatus* sob, AYR04656.1. Clade credibility values are bootstraps from 1000 replicates.

